# MicroRNA and Gene Expression Analysis on Placenta Tissue: An Approach to Understanding Obstetric Antiphospholipid Syndrome at the Molecular Level

**DOI:** 10.1101/780114

**Authors:** Muhammad Aliff Mohamad, Nur Fariha Mohd Manzor, Muhammad Shazwan Suhiman, Jameela Sathar, Hayati Abdul Rahman, Maiza Masri, Nur Syahrina Abdul Rahim, Nazefah Abdul Hamid, Nor Nadeeya Mohamad, Asral Wirda Ahmad Asnawi

## Abstract

Obstetric antiphospholipid syndrome is initiated by the action of antiphospholipid antibodies on placenta. The characteristics of APS in pregnancy include vascular thrombosis, inflammation and impairment of trophoblast implantation. MicroRNA (miRNA) expression has been suggested as one of the genetic factors that contribute to the development of this syndrome. miRNAs regulate gene expressions in a vast assortment of cellular biological mechanisms include the development of placental tissue. Hence, further investigation on the regulation of placental miRNA in APS is required. In this study, we aimed to profile miRNA expressions from placenta tissue of patients with APS. Differentially expressed miRNAs were determined for its targeted genes and pathways. Agilent microarray platform was used to measure placental microRNA expressions between normal placental tissue and those obtained from patients with APS. Differentially expressed miRNAs were detected using GeneSpring GX software 14.2 and sequences were mapped using TargetScan software to generate the predicted target genes. Pathway analysis for the genes was then performed on PANTHER and REACTOME software. Selected miRNAs and their associated genes of interest were validated using qPCR. Microarray findings revealed, 9 downregulated and 21 upregulated miRNAs expressed in placenta of patients with APS. Quantitative expressions of 3 selected miRNAs were in agreement with the microarray findings, however only miR-525-5p expression was statistically significant. Pathway analysis revealed that the targeted genes of differentially expressed miRNAs were involved in several hypothesized signalling pathways such as the vascular endothelial (VE) growth factor (VEGF) and inflammatory pathways. VE-cadherin, ras homolog member A (RHOA) and tyrosine kinase receptor (KIT) showed significant downregulation from the qPCR data while retinoblastoma gene (RET), dual specificity protein phosphatase 10 (DUSP10) and B-lymphocyte kinase (BLK) were significantly upregulated. These preliminary findings suggest the involvement of miRNAs and identified novel associated genes involvement in the mechanism of obstetric APS, particularly through the alteration of vascular-associated regulators and the inflammatory signalling cascade.

## 1.0 Introduction

Based on the Sapporo criteria, Antiphospholipid Syndrome (APS) is characterised by one or more thrombotic or pregnancy related clinical features with the presence of antiphospholipid antibodies [1]. Patients suspected with obstetric APS requires multiple testing at different time points to increase diagnostic precision using current serologic methods. About 70% of successful pregnancies brought to term have been achieved in obstetric APS cases with the use of early and accurate diagnosis which has enabled prompt initiation of appropriate treatment [2]. Nevertheless, there are still another 30% of women at risk of unsuccessful pregnancies amongst obstetric APS patients even when aided with careful treatment and medication [2]. This phenomenon may suggest the existence of other mechanism of obstetric APS. In addition, a number of studies have indicated the importance to review the diagnostic strategy of APS as devastating clinical manifestations have been demonstrated in spite of consistently negative laboratory results [3]

In earlier reports, obstetric APS was associated with placental infarction or placental thrombosis [4]. This finding suggests the importance of placenta in the pathogenesis of obstetric APS. The closest study to APS in relation to this mechanism is related to preeclampsia, where it was revealed that a small regulatory molecule known as ‘microRNA’ (miRNA) was involved in the pathogenesis of preeclampsia [5]. miRNA, is a short sequence of 22 nucleotides non-coding RNAs, may interfere with post-transcriptional gene regulation. This interference could result in inhibition of mRNA translation and mRNA degradation. Until now, about 30% of genes are reported to be regulated by miRNAs [6].

It has been shown that miRNAs could have many target genes and one gene can be targeted by multiple miRNAs [7]. Hence, this gives rise to an interesting avenue on how miRNA-genes network interacts in underlying molecular mechanism of obstetric APS. Recently, there have been many studies suggesting that miRNAs are essential regulators of placental development. miRNAs have been shown to be involved in many crucial processes such as cell differentiation, apoptosis, inflammation and angiogenesis [8]. Furthermore, aberrant miRNA expression has been associated with pregnancy complications and APS. miRNA has also been reported to affect endothelium in recurrent miscarriages [9]. Abnormal expression of miRNAs in disease condition may be used as an indicator of diseases, serve as potential biomarker and therapeutic target [10]. As miRNAs have been proven to involve various mechanisms of disease related to the placenta, this study was done to explore miRNA expression in placenta tissue of patients with obstetric APS. Further analysis on the miRNAs’ targeted genes and pathways were also performed to reveal the possible biomarkers of obstetric APS and its signalling.

## 2.0 Methodology

This is comparative cross-sectional study involving patients diagnosed with obstetric APS. Samples consisted of placenta tissues or aborted product of conceptus and peripheral blood samples which were obtained from consenting patients that experienced a miscarriage, preeclampsia and intrauterine death according to the APS standardised Sydney criteria. A detailed inclusion and exclusion criteria for sample recruitment (APS and normal control group) are summarised in Table 1.

**Table 1.** Inclusion and exclusion criteria for sample selection

| Normal group |  |
| --- | --- |
| Inclusion criteria | Exclusion criteria |
| • Normal spontaneous vaginal delivery or through caesarean procedure.<br>• Does not have pregnancy loss history.<br>• Age ranging from 18-40 years' old<br>• Negative for all LA-aPTT, dRVVT and anti-β2GP1 test | • Intrauterine infectious diseases such as Toxoplasmosis, Other infections (Coxsackievirus, Chicken Pox, Parvovirus B19, Chlamydia, HIV, Syphilis, Human T-lymphotropic Virus), Rubella, Cytomegalovirus and Herpes Simplex virus-2 or<br>• Gestational Diabetes Mellitus, or<br>• Cervical Incompetence, or<br>• Related Autoimmune Disease such as Systemic Lupus Erythematosus.<br>• Positive for LA-aPTT and/or dRVVT and/or anti-β2GP1 test |

| APS group |  |
| --- | --- |
| Inclusion criteria | Exclusion criteria |
| • Successful birth where mother is already diagnosed with aPLs antibodies<br>• History of recurrent miscarriages ($\geq 3$) at $\geq 16$ weeks' period of gestation.<br>• History of 2 miscarriages, current miscarriage at $\geq 16$ weeks' period of gestation.<br>• Complication of intrauterine death<br>• Delivery with preeclampsia or severe eclampsia at any gestation period.<br>• Intrauterine Growth Restriction at any gestation period.<br>• Age ranging from 20 to 40 years' old<br>• Positive for LA-aPTT and/or dRVVT and/or anti-$\beta 2$GP1 test | • Intrauterine infectious diseases such as Toxoplasmosis, Other infections (Coxsackievirus, Chicken Pox, Parvovirus B19, Chlamydia, HIV, Syphilis, Human T-lymphotrophic Virus), Rubella, Cytomegalovirus and Herpes Simplex virus-2 or<br>• Gestational Diabetes Mellitus, or<br>• Cervical Incompetence, or<br>• Related Autoimmune Disease such as Systemic Lupus Erythematosus.<br>• Negative for all LA-aPTT, dRVVT and anti-$\beta 2$GP1 test |

### 2.1 Sample collection

Samples were immersed in 3-5 ml RNA later (Ambion™, USA) in 50 ml tubes and stored in - 80°C within 24 hours until further process of RNA extraction. About 9 ml whole blood were collected in citrated tube (BD Vacutainer, USA) for lupus anticoagulant, anticardiolipin and anti-B2GP1 antibody testing by certified haematology laboratory.

### 2.2 Total RNA extraction, purification and quality control

Total RNA was extracted based on procedure provided by PureLink RNA Mini Kit (Ambion™, USA). RNA pellet was then purified and measured for its quality. Sample with the reading within ~2.0 for both A260/280 and A260/230 were selected for further RNA Integrity Number (RIN) measurement. This step was performed using Agilent 2100 Bioanalyzer (Agilent, USA). A minimum threshold of RIN ~7.0 was set to eliminate experimental bias of poor RNA quality control prior to microarray profiling quantification. Samples that passed the quality-check were stored in −80°C for microarray analysis and quantitative polymerase chain reaction (qPCR).

### 2.3 MicroRNA microarray analysis

miRNA microarray profiling was conducted using G3 Human miRNA Microarray, Release 21, 8×60k slide (Agilent, USA) following manufacturer’s guideline. It contained 2549 human probes that against human microRNA sequence from miRBase 21.0. Briefly the steps start with preparation of Spike-In solutions, de-phosphorylation, denaturation and ligation of samples followed by drying, hybridization, wash and slides scanning.

From the microarray generated scan image, probe features were extracted by scanning the data with Feature Extraction Software Version 9.5.3. Data gathered from images scanning was then analysed with Gene Spring GX 14.2 software, where the expression value and differentially expressed miRNA data were obtained.

### 2.4 MicroRNA-target genes and pathway analysis

List of differentially expressed miRNA was run in Target Scan program to obtain the target genes of the miRNAs. Fold change of ≥ 2 and P < 0.05 (by modified T-test) was set to filter out the predicted target genes. Next, a set of genes that remained after fold change and p-value filtering was analysed with PANTHER [11] and Reactome program [12] to find the pathway associated with the study. Selected genes from the pathway of interest were further validated using qPCR.

### 2.5 Validation of MicroRNA expression by qPCR

#### 2.5.1 cDNA synthesis for miRNA

miRNA cDNA synthesis was performed using reagents from miScript PCR starter kit (Qiagen, USA). The master mix was prepared in reaction tube (Applied Biosystem) as stated in Table 2. The tube was centrifuged using mini-microcentrifuge (Eppendorf, Germany) and inserted into Mastercycler EP-Gradient Thermal Cycler 96 well (Eppendorf, Germany). Reverse Transcriptase Profile was adjusted according to Table 3.

**Table 2.** cDNA synthesis component for miRNA

| Component | Volume for one tube |
| --- | --- |
| 5x miScript Hiflex buffer | 4 ul |
| 10x miScript nucleomix | 2 ul |
| RNAse free water | 10 ul |
| miScript RT mix | 2 ul |
| Total RNA | 2 ul |
| Total volume | 20 ul |

**Table 3.**
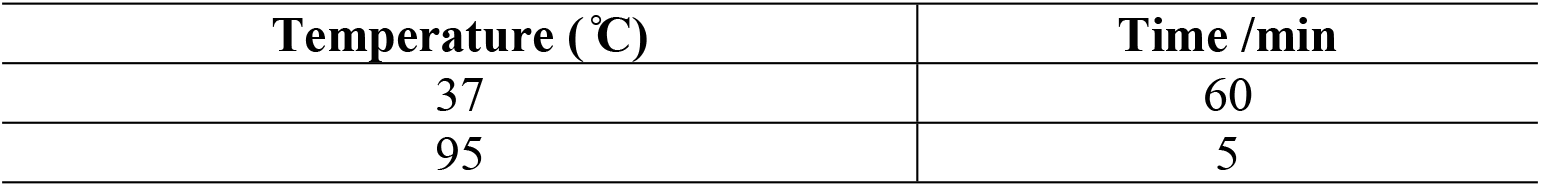
miRNA cDNA cycling program

#### 2.5.2 Primer for miRNA qPCR

Primers for human hsa-miR-3148, hsa-miR-670, hsa-miR-525, RNU6-2 and SNORD48 were purchased from Qiagen, USA.

#### 2.5.3 Preparation of master mix and qPCR protocol for miRNA quantification

Reagents from MiScript PCR starter kit (Qiagen, USA) were used for the preparation of master mix for miRNA qPCR. The master mix components were prepared in MicroAmp fast reaction tube (Applied Biosystem) as stated in Table 4. Quantitative PCR (qPCR) for miRNA was performed using Step OnePlus Real Time PCR System (Applied Biosystem, USA). Prior to starting the protocol, the qPCR machine was checked to ensure its connection with the Step OnePlus Software Ver 2.3 and programmed according to the profile as written in Table 5.

**Table 4.**
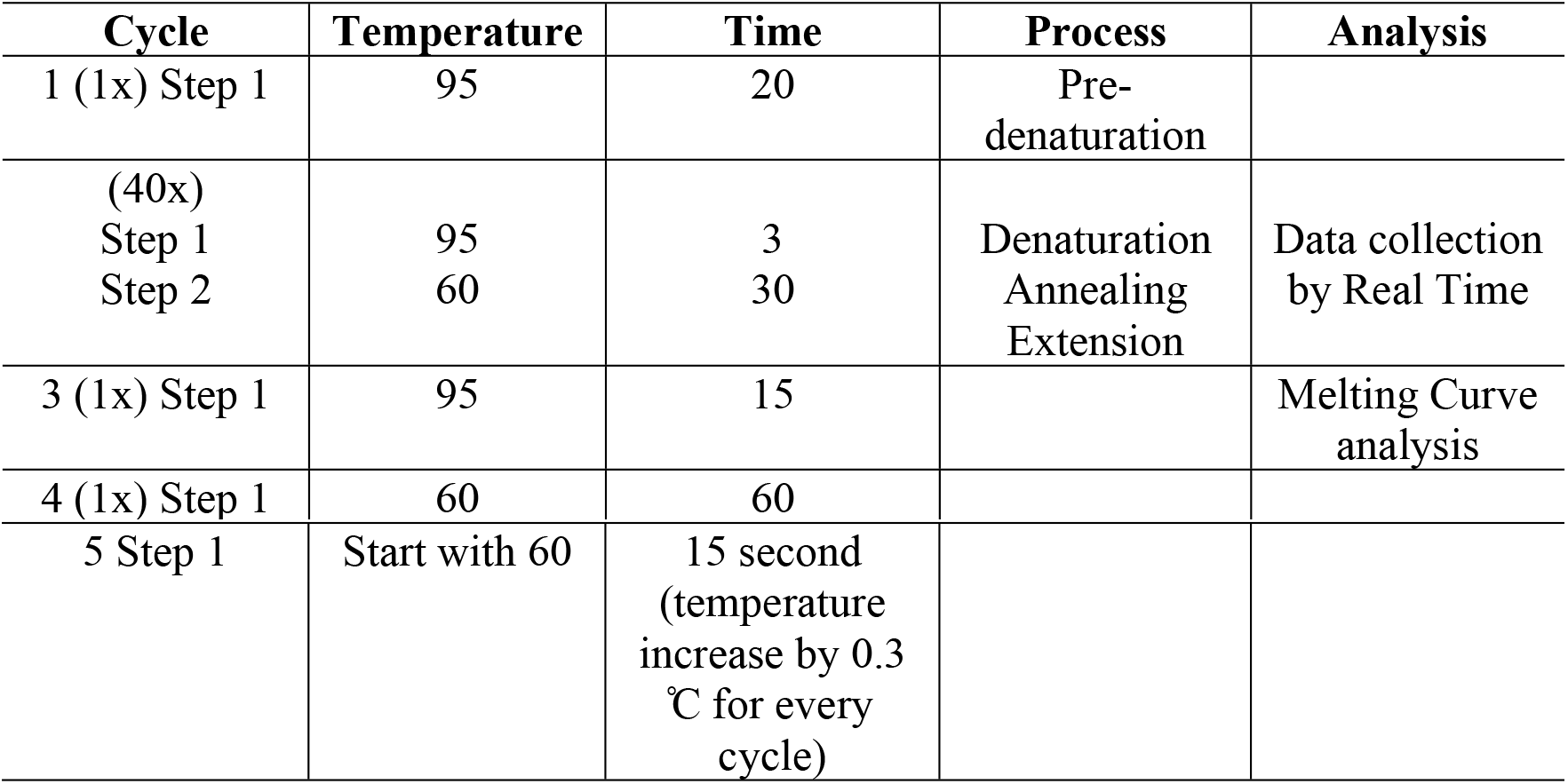
Master mix components for miRNA qPCR

| Cycle | Temperature | Time | Process | Analysis |
| --- | --- | --- | --- | --- |
| 1 (1x) Step 1 | 95 | 20 | Pre-denaturation |  |
| (40x)<br>Step 1<br>Step 2 | 95<br>60 | 3<br>30 | Denaturation<br>Annealing<br>Extension | Data collection<br>by Real Time |
| 3 (1x) Step 1 | 95 | 15 |  | Melting Curve<br>analysis |
| 4 (1x) Step 1 | 60 | 60 |  |  |
| 5 Step 1 | Start with 60 | 15 second<br>(temperature<br>increase by 0.3<br>°C for every<br>cycle) |  |  |

**Table 5.** Cycling program for miRNA qPCR

| Component | Volume for one tube |
| --- | --- |
| 2x QuantiTect SYBR Green PCR Master Mix | 12.5 ul |
| 10x miScript Universal Primer | 2.5 ul |
| 10x miScript primer assay | 2.5 ul |
| RNase free water | 2.5 ul |
| Template cDNA | 7.0 ul |
| Total volume | 25 ul |

### 2.6 miRNA targeted-gene expression analysis by qPCR

#### 2.6.1 cDNA synthesis for targeted-genes

For cDNA synthesis of targeted-genes, Superscript III First-Strand Synthesis SuperMix (Invitrogen, USA) was used. Reaction mix and RT profile were detailed out as in Table 6 and 7 respectively.

**Table 6.** Component mixture of master mix for targeted-genes cDNA synthesis

| Components | Volume in one tube |
| --- | --- |
| 2x reaction mix | 10.0 ul |
| DEPC-treated water | 6.0 ul |
| Total RNA | 2.0 ul |
| RT enzyme | 2.0 ul |
| Total final volume | 20 ul |

**Table 7.**
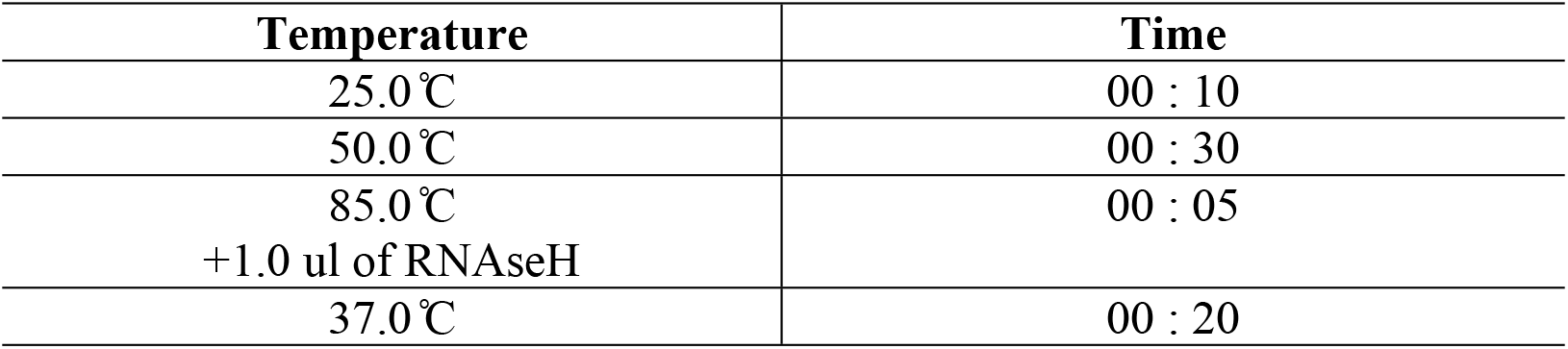
Invitrogen cDNA cycling program for targeted-genes

#### 2.6.2 Primer sequences for qPCR

The primer sequences were designed using primer 3 input (http://primer3.ut.ee) based on the sequences of the coding region of each gene as referred to the Gene bank data. All new primers are listed in Table 8 and 9.

**Table 8.**
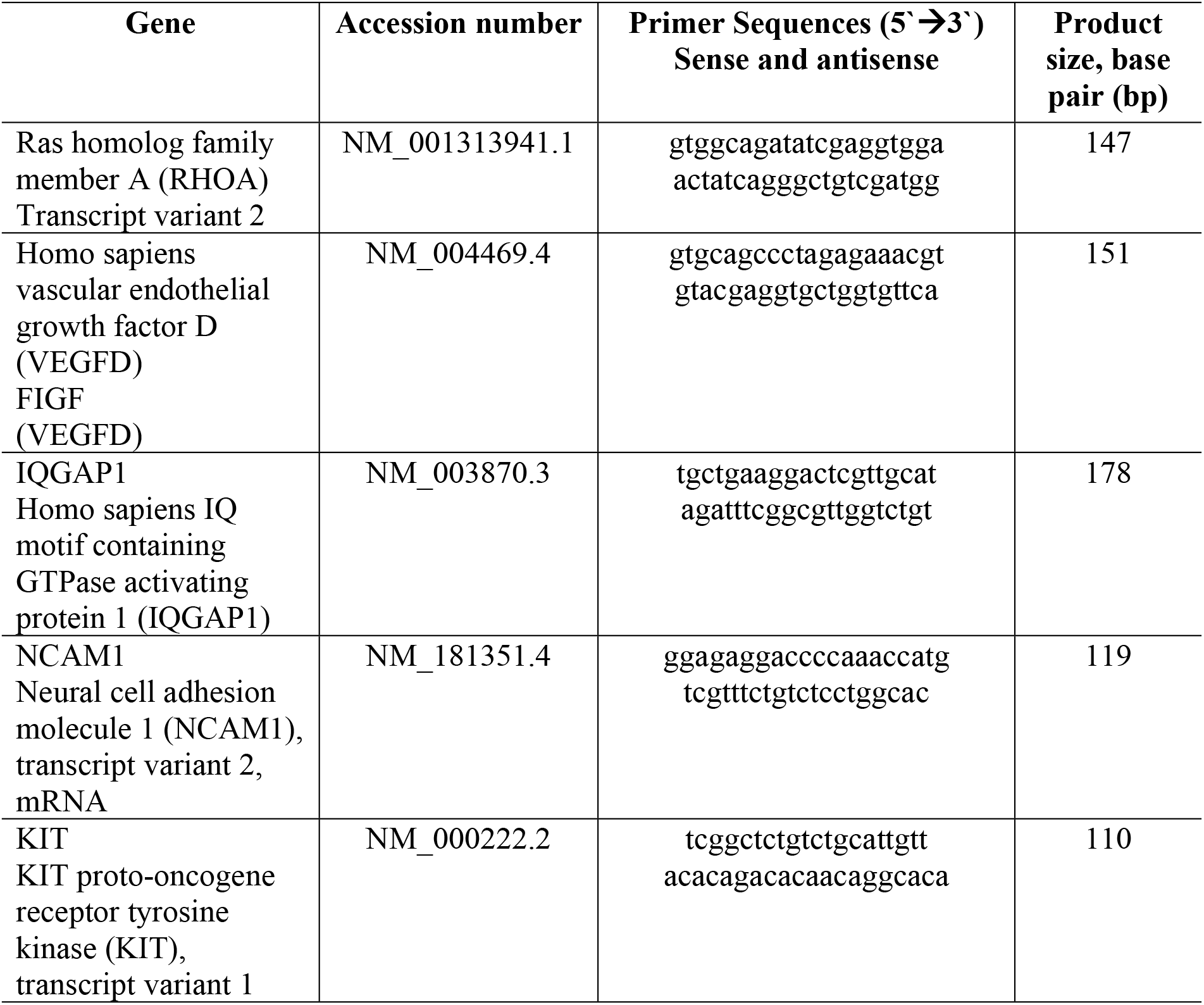
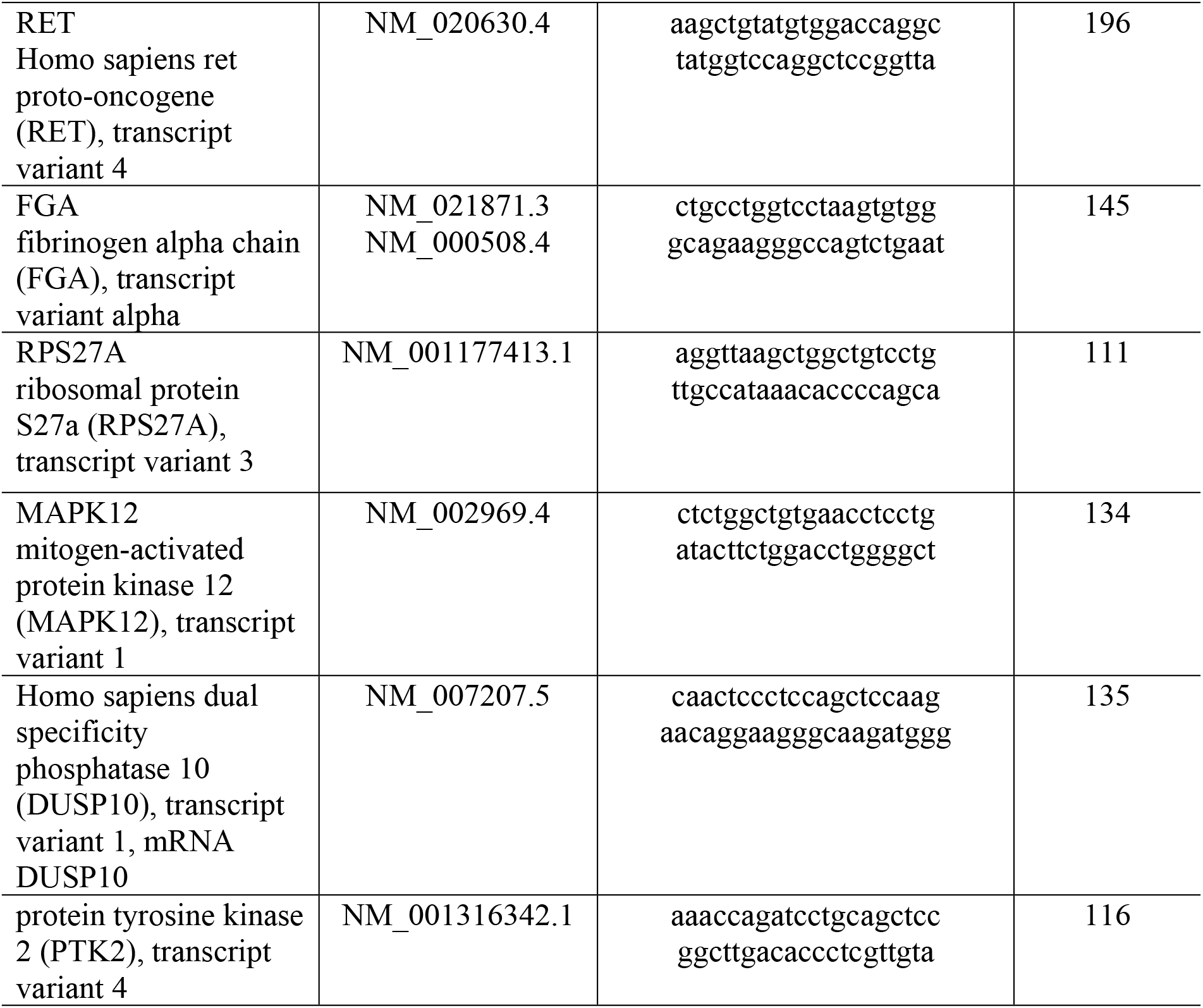
Primer sequences for upregulated miRNA targeted-genes

**Table 9.** Primer sequences for downregulated miRNA targeted-genes

| **Gene** | **Accession**<br>**Number** | **Primer Sequences (5'-3')**<br>**Sense and anti-sense** | **Product**<br>**size base**<br>**pair (bp)** |
| --- | --- | --- | --- |
| B-Lymphocyte Kinase<br>(BLK) | NM_001715.2 | ggc cat taa gac gct gaa gg<br>atc ctc tgg cca tgt act cg | 158 |
| Cluster of<br>Differentiation<br>(CD19) | NM_001178098.1 | gaa agc gaa tga ctg acc cc<br>gct gct cgg gtt tcc ata ag | 182 |
| Mitogen-Activated<br>Protein Kinase-Activated Protein<br>Kinase 2<br>(MAPKAPK2) | NM_004759.4 | acc gta cta tgt ggc tcc ag<br>atg gtc att ctc tgg gtg gg | 270 |
| Myosin Heavy Chain<br>15 (MYH15) | NM_014981.1 | gta gat gac ctc ctg acc cg<br>ctt ttg tgc tgc cag gtc at | 150 |

Five genes; Glyceraldehyde 3-phosphate dehydrogenase (GAPDH), Platelet and endothelial cell adhesion molecule-1 (PECAM-1), Vascular endothelial cadherin (VE-cadherin), Endothelial Nitric Oxide Synthase (eNOS) and Von Willebrand factor (vWF) were selected from angiogenic-endogenic-associated genes studied by Nur Fariha et al (2012). These four genes were selected based on its association to the pathway analysis of upregulated miRNAs-targeted genes.

#### 2.6.3 Preparation of qPCR master mix and cycling protocol for targeted-genes quantification

Mastermix solution was prepared by using SsoAdvance Universal SYBR Green Supermix kit (BioRad, USA). Reaction mix was prepared as according to Table 10 and qPCR was performed using Step OnePlus Real Time PCR System (Applied Biosystem, USA) according to the profile stated in Table 11.

**Table 10.** Master mix components for targeted-genes qPCR

| Component | Volume for one tube |
| --- | --- |
| RNase DNase distilled water | 7 ul |
| 2x Sso Advance Universal SYBR Green Supermix | 10 ul |
| cDNA | 1 ul |
| Forward Primer | 1 ul |
| Reverse Primer | 1 ul |
| Total volume | 20 ul |

**Table 11.**
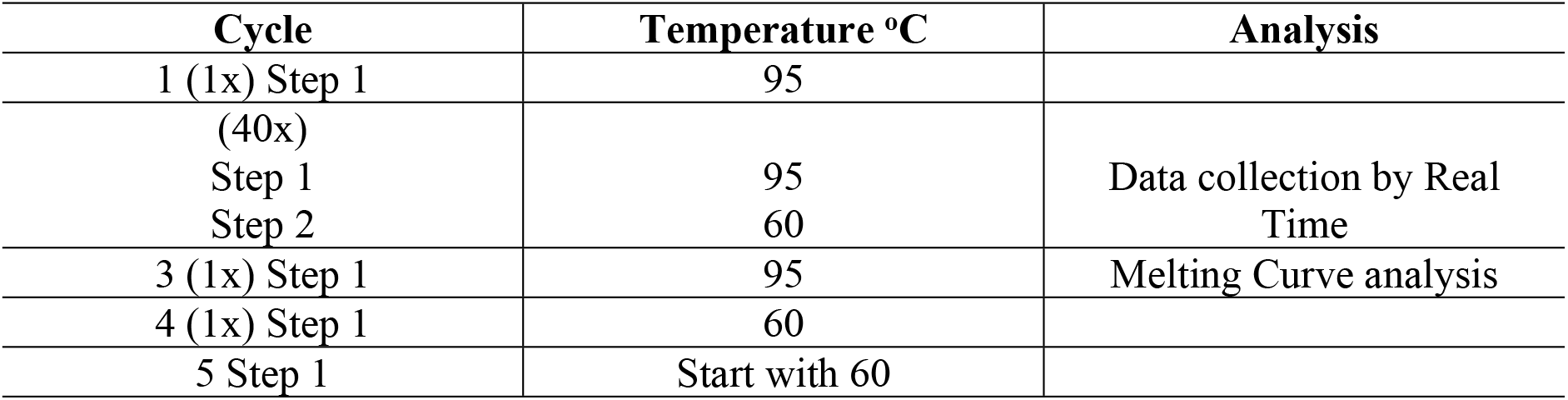
qPCR cycling program for targeted-genes quantification

### 2.7 Statistical analysis

The statistical analysis was performed using GraphPad Prism Software v5.0. The qRT-PCR results are expressed as mean + standard deviation (SD) and statistically analysed using Mann Whitney test for comparison of 2 groups, with p < 0.05 considered significant was used to determine significance of genes and miRNAs expression between APS and normal samples.

## 3.0 Result

### 3.1 Patient’s data profile

Placenta tissue and blood sample from 5 normal and 3 APS cases were subjected to serological test and microRNA microarray analysis. All samples from normal cases were negative for LA, aCL and anti-β2GP1 tests. Pregnancy complications and serological test results for all 3 APS cases were detailed out in Table 12.

**Table 12.** Data profile of APS sample

| Sample | Complications | Tests |  |  |
| --- | --- | --- | --- | --- |
|  |  | LA<sup>a</sup> | aCL<sup>b</sup> | anti-β2GP1<sup>c</sup> |
| APS 1 | Early miscarriage | Positive | Positive | Negative |
| APS 2 | PE, miscarriage | Positive | Positive | Positive |
| APS 3 | Early miscarriage | Positive | Positive | Negative |

<sup>a</sup>LA, Lupus anticoagulant; <sup>b</sup>aCL, anticardiolipin; <sup>c</sup>anti-β2 Glycoprotein 1

### 3.2 MicroRNA microarray profile

#### 3.2.1 Volcano plot, Principal component analysis (PCA) plot and Hierarchical clustering of samples

Through filtering on volcano plot with corrected *p*-value cut-off value of 0.05 (*p* < 0.05) and fold change cut-off value of 2.0 (>2.0), 30 miRNAs were detected as differentially expressed; 21 were upregulated and 9 were downregulated as shown in Fig 1 (A). From the figure, miRNA that are highly upregulated are localized farther to the right side and miRNA that are highly downregulated are farther to the left side. Upregulated miRNAs with statistical significance less than 0.05 were indicated in red whereas downregulated miRNAs with statistical significance less than 0.05 was indicated in blue. miRNAs which expressed fold change less than 2 and not statistically significant appear in grey. Differentially expressed miRNAs are listed in Table 13. An exploratory PCA was performed using the analysis platform embedded in the GeneSpring GX 14.5 Software. PCA plot in Fig 1 (B) shows the distinct arrangements of samples between normal and APS groups. In this study, hierarchical clustering was performed on a one-channel microarray data. The dendrogram provided some qualitative means of assessing the similarity or differences of miRNAs profiling in patient samples. Dendrogram of comparison between normal versus APS is shown in Fig 2. From this heat map, technical and biological replicates of the same group of samples were grouped together.

**Fig 1.**
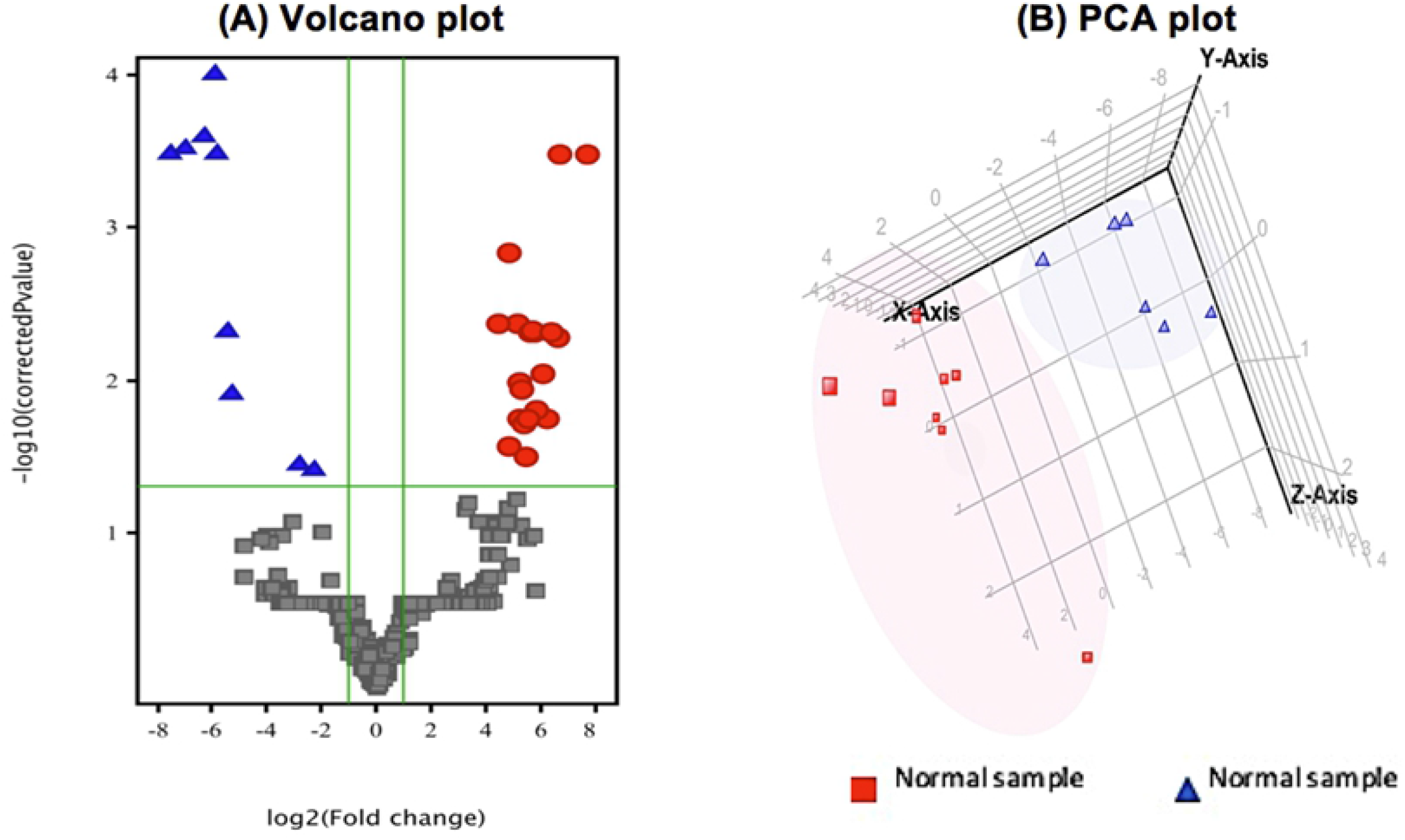
Volcano and PCA plot for comparison between APS and normal samples. Volcano (A) and PCA (B) plots for APS versus normal samples. For the volcano plot, up- and downregulated miRNA were indicated in red and blue respectively. whereas downregulated miRNA. miRNAs which appear in grey were not statistically significant in expression.

**Table 13.** Differentially expressed miRNAs in normal versus APS samples

| Upregulated miRNAs | Downregulated miRNAs |
| --- | --- |
| hsa-miR-1238-5p | hsa-miR-1183 |
| hsa-miR-1825 | hsa-miR-1914-3p |
| hsa-miR-2278 | hsa-miR-4497 |
| hsa-miR-3148 | hsa-miR-4689 |
| hsa-miR-3907 | hsa-miR-518e-5p |
| hsa-miR-3912-5p | hsa-miR-525-5p |
| hsa-miR-4436b-5p | hsa-miR-564 |
| hsa-miR-4502 | hsa-miR-6768-5p |
| hsa-miR-4644 |  |
| hsa-miR-4730 |  |
| hsa-miR-4769-3p |  |
| hsa-miR-5007-5p |  |
| hsa-miR-595 |  |
| hsa-miR-670-5p |  |
| hsa-miR-6739-5p |  |
| hsa-miR-6751-3p |  |
| hsa-miR-6751-5p |  |
| hsa-miR-6775-3p |  |
| hsa-miR-6876-5p |  |
| hsa-miR-7844-5p |  |
| hsa-miR-7850-5p |  |

**Fig 2.**
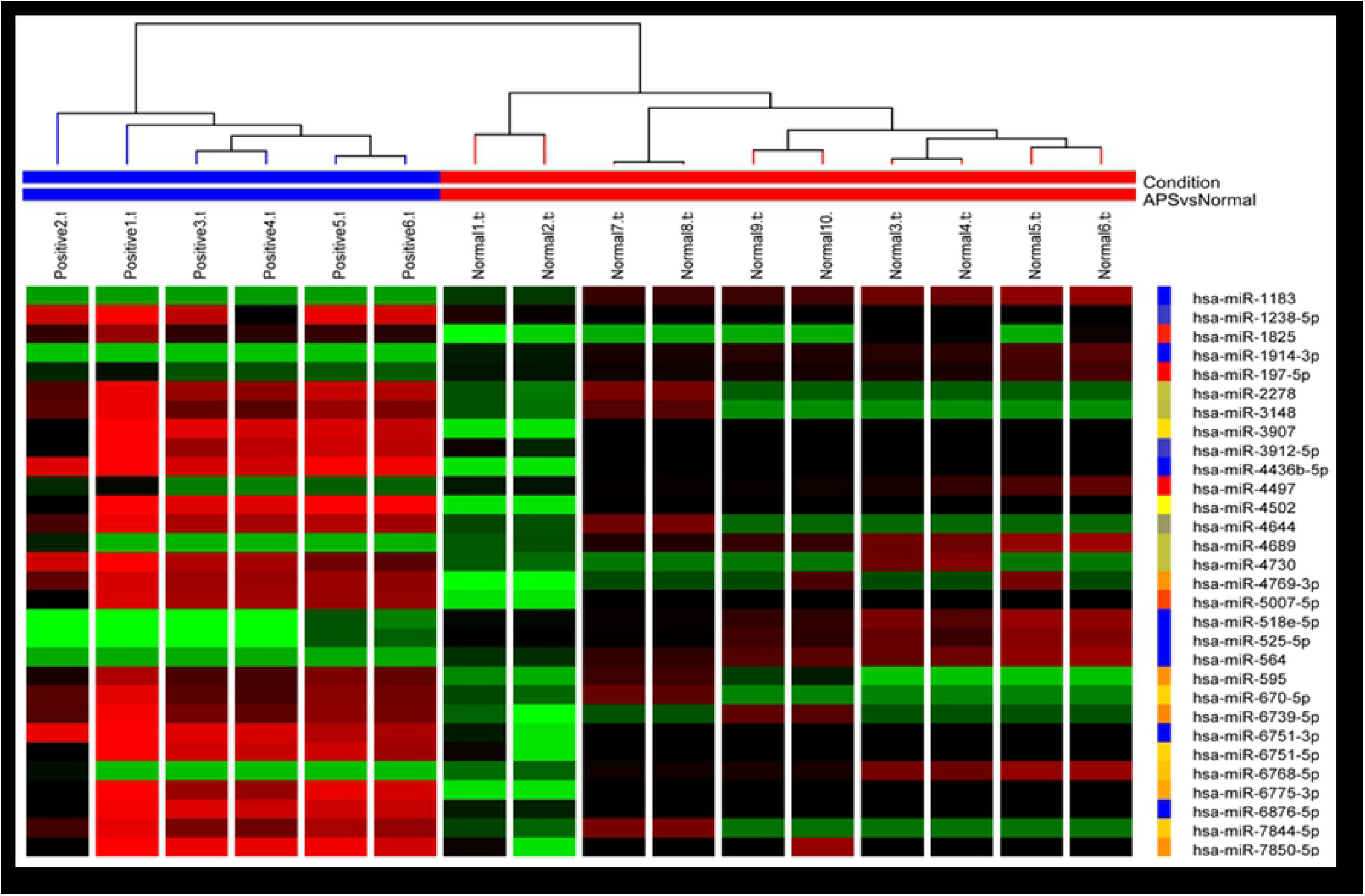
Hierarchical clustering of normal versus APS samples. The colours represent the expression levels of the miRNAs. Red represent high expression while green represent low expression. The right ledger shows the differentially expressed miRNAs and top ledger shows the samples.

### 3.3 MicroRNA-targeted genes and pathways

When comparing normal and APS samples using the computational algorithm of TargetScan, it was predicted that the upregulated and downregulated miRNAs would target 578 and 178 genes respectively. Gene ontology (GO) analysis of the listed genes were performed through DAVID, specifically on the Biological Process (BP) GO term. Only the top 10 BP GO term were listed in Table 14 and 15.

**Table 14.** GO analysis on biological process of upregulated miRNAs-targeted genes

| GO Term | Gene count | p-value |
| --- | --- | --- |
| endoplasmic reticulum mannose trimming | 11 | 6.6E-3 |
| G2/M transition of mitotic cell cycle | 4 | 1.2E-2 |
| response to fatty acid | 4 | 1.4E-2 |
| response to vitamin A | 5 | 1.5E-2 |
| glucose transport | 4 | 1.7E-2 |
| negative regulation of endoplasmic reticulum stress-induced intrinsic apoptotic signaling pathway | 4 | 1.9E-2 |
| flavonoid biosynthetic process | 4 | 2.5E-2 |
| flavonoid glucuronidation | 11 | 2.5E-2 |
| metabolic process | 3 | 3.2E-2 |

**Table 15.** GO analysis on biological process of downregulated miRNAs-targeted genes

| GO Term | Gene Count | p-value |
| --- | --- | --- |
| Response to nutrient | 15 | 6.0E-3 |
| Notch signalling pathway | 14 | 9.3E-3 |
| Optic nerve development | 4 | 9.4E-3 |
| G2/M transition of mitotic cell cycle | 14 | 2.4E-2 |
| Carboxylic acid metabolic process | 44 | 2.4E-2 |
| Oxoacid metabolic process | 44 | 2.6E-2 |
| Positive regulation of sperm motility | 3 | 2.7E-2 |
| Atrioventricular valve formation | 3 | 2.7E-2 |
| Kidney development | 18 | 2.8E-2 |
| Response to vitamin A | 4 | 2.9E-2 |

Further pathway analysis of upregulated miRNAs resulted in 1168 pathways. Considering that the mechanism of obstetric APS is through the alteration of vascular functions, VEGF-VEGFR pathway was selected as the main focus for further gene validation. Meanwhile, 71 pathways were found to involve the targeted genes of downregulated miRNAs. Among all, a few pathways of interest related to inflammatory processes were noted. The pathways include B-cell activation, interleukin signalling pathway, inflammation mediated by chemokine and cytokine signalling pathways and toll receptor signalling pathways. Several genes which were chosen for qPCR analysis from the selected pathways were listed in Table 16.

**Table 16.**
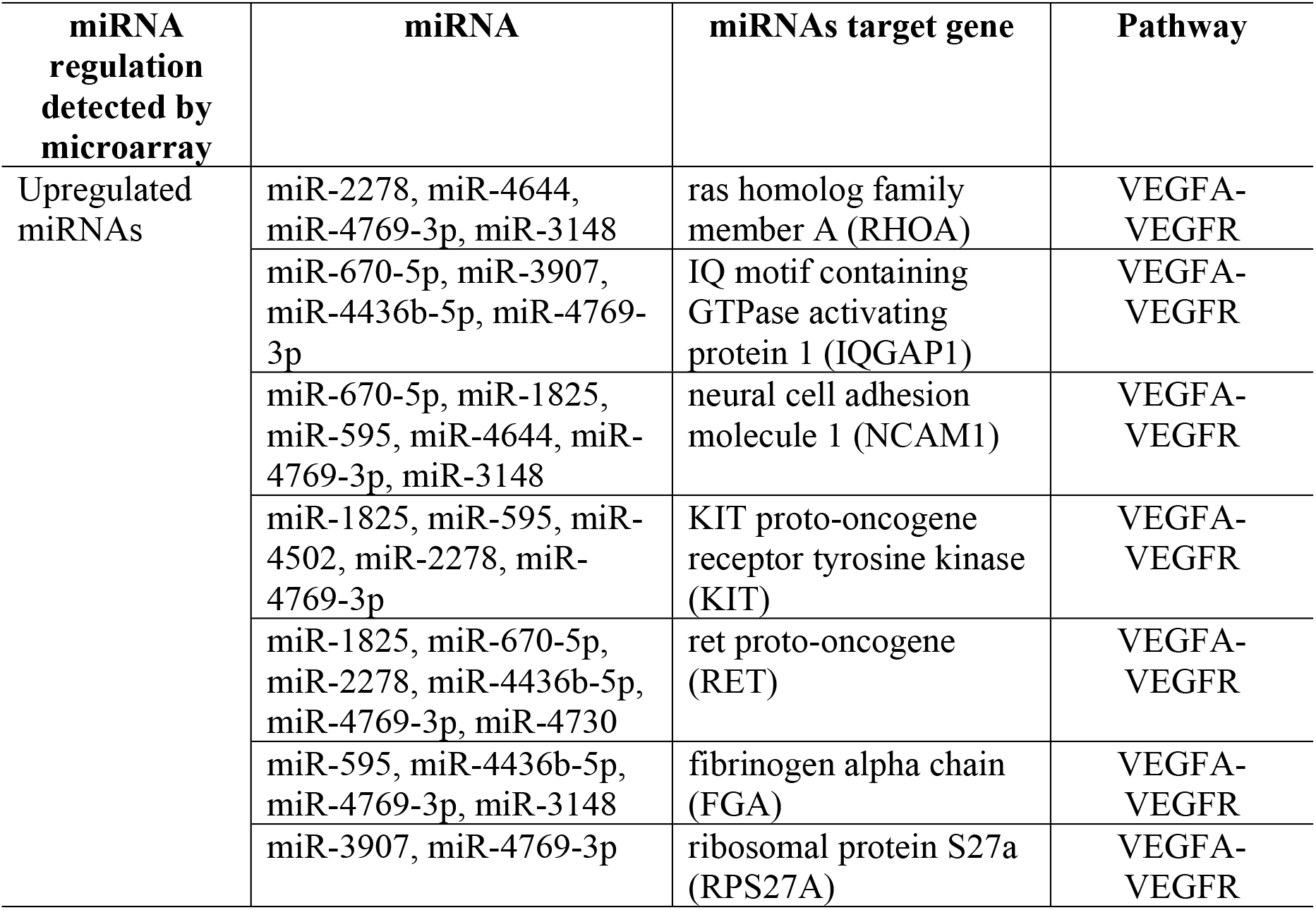
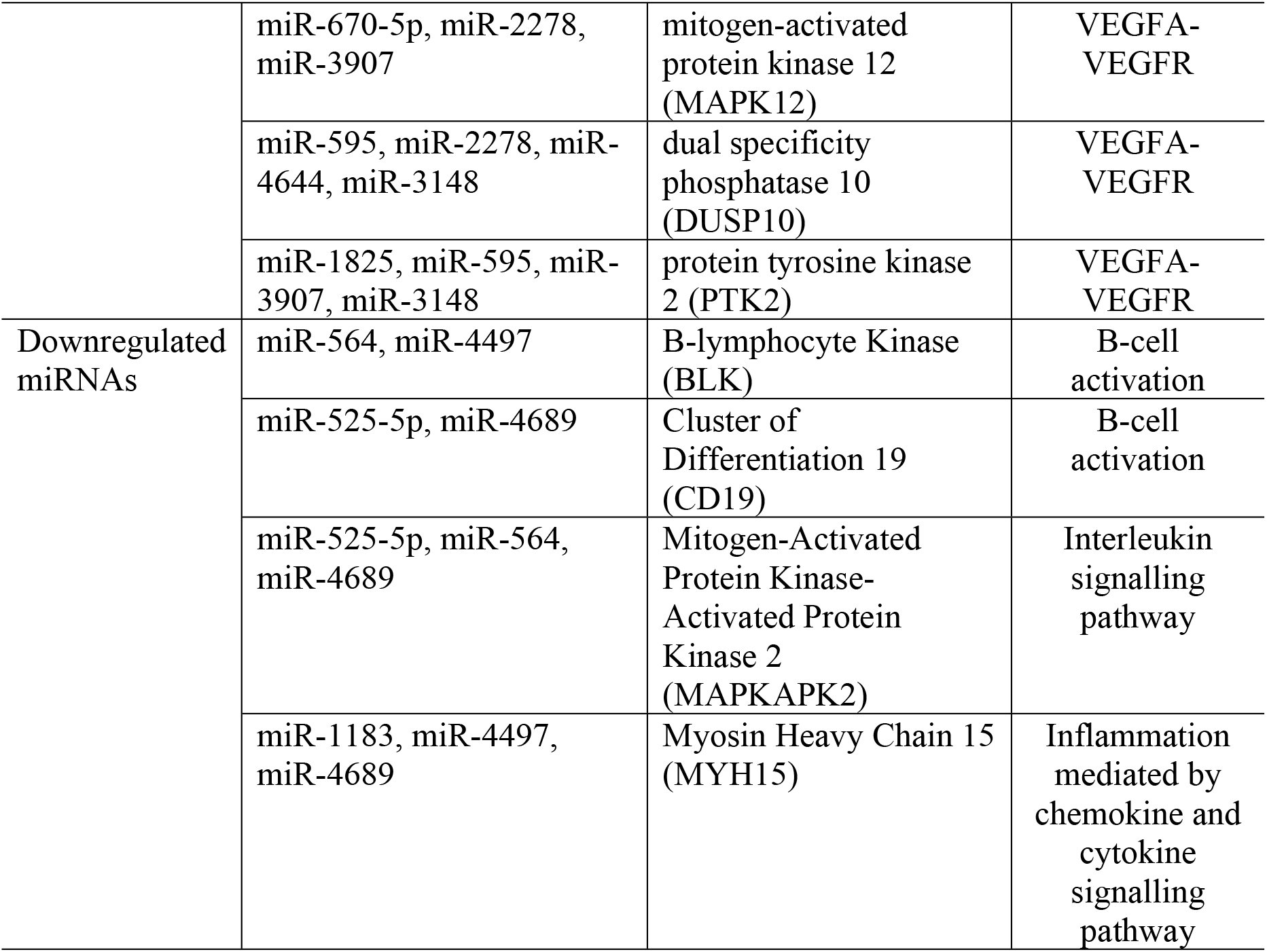
Selected genes from pathways of interest

### 3.4 Quantitative expression of miRNAs and genes

We further quantified the miRNAs; miR-670 and miR-3148 were upregulated in APS as compared to normal by 3 and 4-fold changed respectively. However, both differentially expressed miRNAs were not significant (P < 0.05, Mann-Whitney U-test). Another miRNA, miR-525-5p which was downregulated from the microarray findings were also quantified using qPCR. The results showed a significant downregulation of miR-525-5p in APS group as compared to the normal group (Table 17 and Fig 3)

**Table 17.** Expression pattern of selected miRNAs from qPCR data

| Systematic name | P value | Regulation<br>*APS compare to normal | Fold change |
| --- | --- | --- | --- |
| miR-670 | 0.5167 | UP | 3.94 |
| miR-3148 | 0.4121 | UP | 4.99 |
| miR-525-5p | 0.0121 | DOWN | 232.14 |

**Fig 3.**
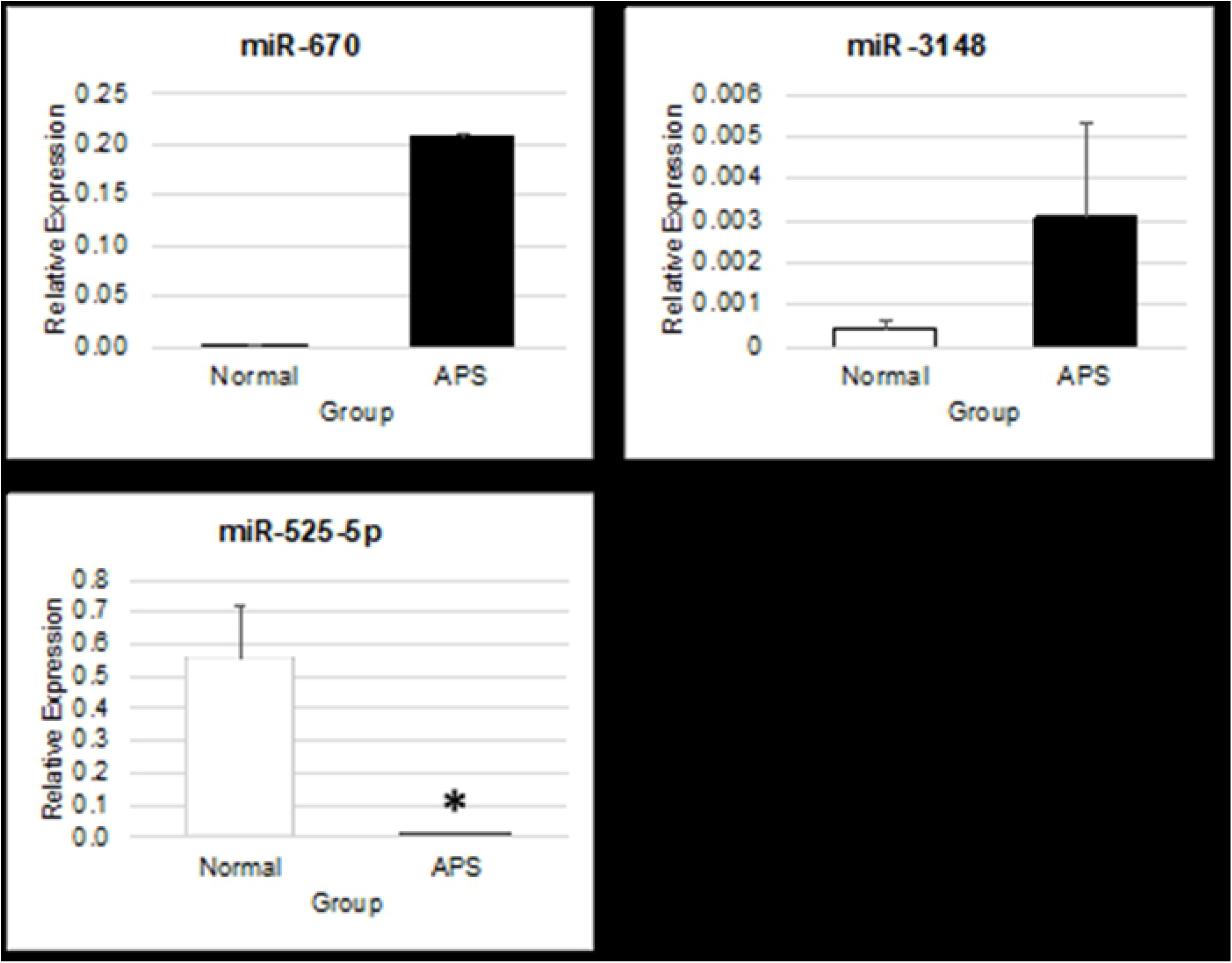
Relative expression of miR-670, miR-3148 and miR-525-5p. Relative expression (mean ± SEM) of miR-670, miR-3148 and miR-525-5p in normal placenta (Normal, n = 4) and placenta of patient with APS (APS, n = 2), determined by real-time PCR. (* indicate p < 0.05)

### 3.5 Quantitative expression of predicted miRNA-targeted genes

#### 3.5.1 Upregulated miRNA-targeted genes

From the qPCR data, most of the genes (10 genes) were upregulated; PECAM-1, vWF, Ve-cadherin, RHOA, IQGAP, NCAM1, KIT, FGA, RPS27A and PTK2, while 3 genes; RET, MAPK12 and DUSP10 were downregulated. On the other hand, eNOS expression showed no difference in APS tissue compared to normal (Fig 4). However only 5 genes, VE-cadherin, RHOA, KIT, RET and DUSP10 showed significant difference compared to normal sample (p < 0.05, Mann-Whitney U test)

**Fig 4.**
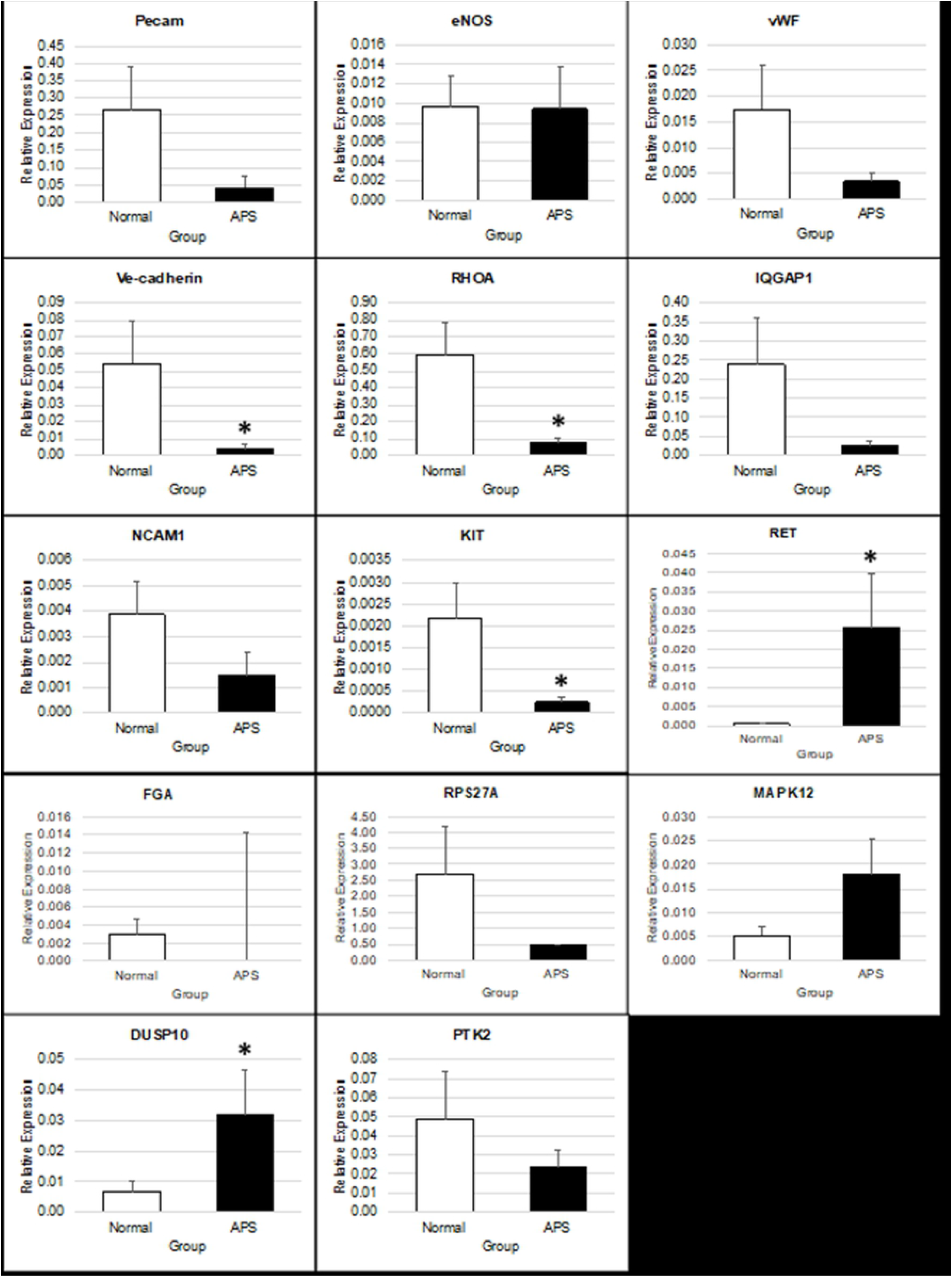
Relative expression of upregulated miRNA-targeted genes. Relative expression (mean ± SEM) of upregulated miRNA-targeted genes and vascular-associated genes in placentas of patient with APS (APS, n = 2) and normal (Normal, n = 4), determined by real-time PCR. (* indicate p < 0.05)

#### 3.5.2 Downregulated miRNA-targeted genes

Based on Fig 5, out of 4 genes quantified by qPCR only BLK was significantly upregulated between the groups

**Fig 5.**
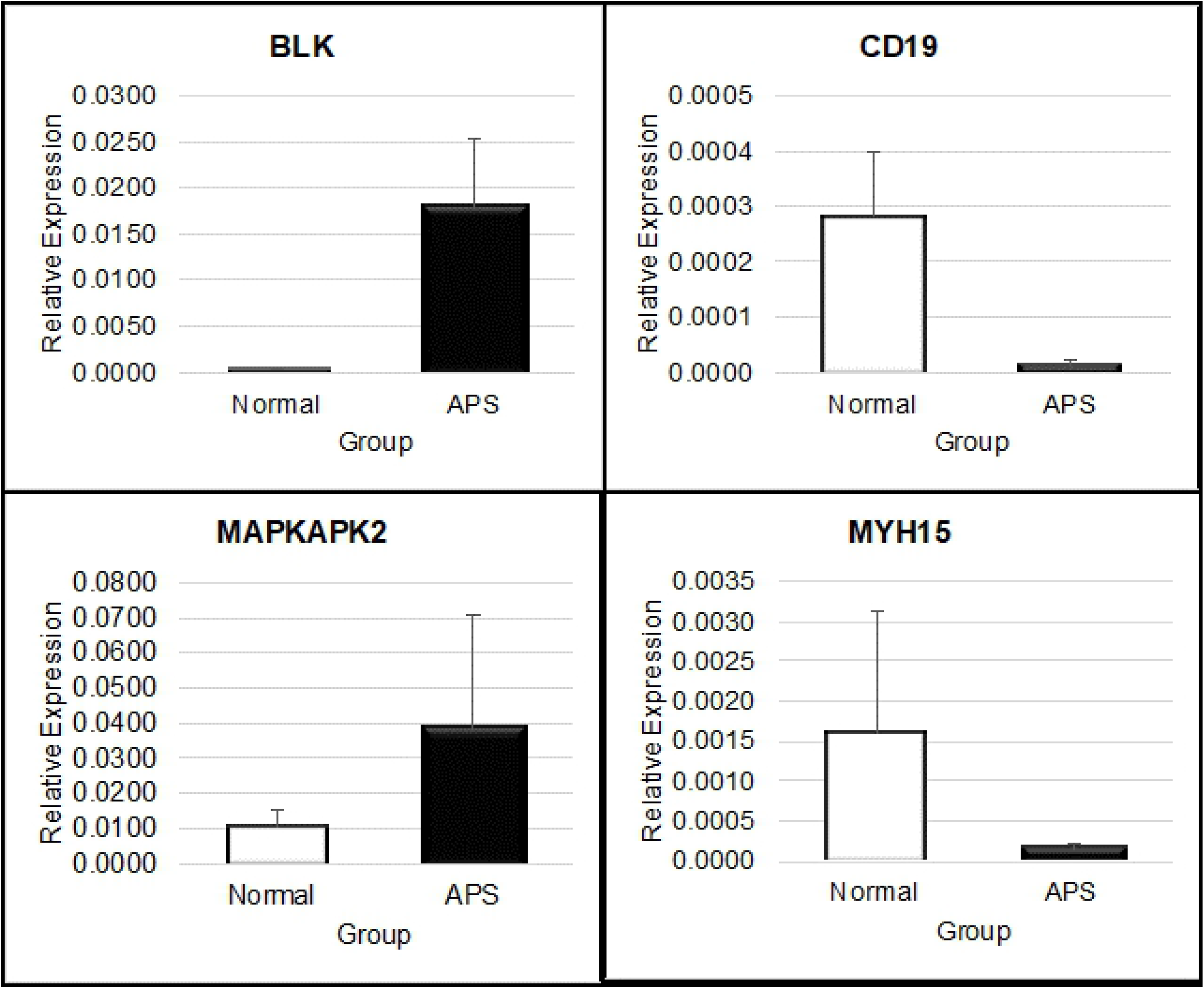
Relative expression of downregulated miRNA-targeted genes. Relative expression (mean ± SEM) of downregulated miRNA-targeted genes in placentas from obstetric APS (APS, n = 2) and normal (Normal, n = 4), determined by real-time PCR. (* indicate p<0.05)

## 4.0 Discussion

The concept of miRNAs-regulated mechanism of disease has long been recognised but the involvement of this molecule in APS remained largely unknown. Prior work has documented the involvement of miRNA in various underlying molecular mechanisms of diseases [13]. Teruel, for example, reports that downregulation of miR-19b and miR-20a which are members of the miR-17~92 cluster was inversely correlated with expression of tissue factor (TF). TF is known as a primary physiological initiator of blood coagulation. Thus, increase TF expression could trigger hypercoagulable state [14]. These studies have mostly been designed based on genes of interest known to be expressed in certain cell types and miRNAs that have been acknowledged as main players in a specific type of cell. In this study, miRNA was chosen through in silico method. This method predicts miRNA using web databases of miRNA’s algorithm. This selection is based on genes of interest which were uploaded in the database. By using microarray analysis, we found miRNAs expression that is unique to APS were detected. From the PCA plot and hierarchical clustering, clear separation between normal and APS groups were observed which revealed a distinct pattern of miRNAs expression in both groups. There were 30 differentially expressed miRNAs identified, where 21 miRNAs were significantly upregulated and 9 miRNAs were downregulated.

From the 21 differentially upregulated miRNAs identified, miRNA-3148 was predicted to target the highest number of genes with 12 genes in total. miR-3148 was studied in autoimmune disease of Systemic Lupus Erythematosus (SLE). Overexpression of miRNA-3148 was found to correlate with the downregulated mRNA expression of Toll-like receptor 7 and vice versa [15]. Meanwhile, miR-670-5p, miR-4644 and miR-595 were shown to target 7 genes. miR-670-5p is associated with cell proliferation that targeted Prospero Homeobox 1 (PROX1). Overexpression of miR-670-5p decreased expression of PROX1 [16]. PROX1 is known as one of the lymphatic markers [17]. Although placenta tissue has no lymphatic vessels, the presence of several numbers of lymphatic markers including PROX1 in placenta has been reported previously, which point out the lymphatic phenotypic characteristics of placental trophoblast [18]. The downregulation of this gene could perturb its role in vessel remodelling and could be linked to vascular thrombosis in placenta of patient with APS. While for miR-595, it was previously found to be overexpressed in the sera of patients with inflammatory bowel disease [19]. As inflammatory-mediated tissue damage is also indicated in placenta of patients with APS, miR-595 could possibly involve in the mechanism of this disease. On the other hand, miR-4644 has been identified as potential diagnostic marker for pancreatic cancer [20]. At the moment, no other diseases have been connected to the overexpression of miR-4644 found in literature.

Interestingly, differential miRNAs in APS identified in our study was distinctly different from non-APS miRNA of placenta from early pregnancy loss and PE [21–23]. When we BLAST the predicted target genes of the upregulated miRNAs using REACTOME platform, the genes were connected to the VEGF-VEGFR pathway. This pathway consists of vascular-associated genes that regulate the angiogenesis process. Angiogenesis is a process of new blood vessels formation from pre-existing vasculature that also happened in wound healing and embryogenesis [24]. A study by Di Simone and co-workers suggested that one of the causes of aPL-mediated foetal loss was inhibition of angiogenic factor secretion and direct impairment of the placenta angiogenesis [25]. The possible effects of aPLs include limiting trophoblast migration [26], suppressing placenta growth factor production [27] and also modulation of trophoblast angiogenic factor secretion [28].

The qPCR findings of genes in VEGF-VEGFR pathway supported our hypothesis that the targeted genes were predicted to be downregulated due to suppression by the upregulated miRNAs. However, it is worth noting there were targeted genes that were either upregulated or unchanged in its expression. VE-cadherin, RHOA and KIT were significantly downregulated while RET and DUSP10 were significantly upregulated (Fig 4). VE-cadherin has been found to enhance cytotrophoblast invasiveness [29]. Damsky et al reported that antiphospholipid antibodies caused the down regulation of VE-cadherin, hence causing impaired trophoblast invasion [30]. Meanwhile, RHOA, which is known as a modulator of gene expression, adhesion and migration of activated macrophage, plays an important role in NFκB inflammatory signalling [31]. RHOA protein was found to be significantly increased in monocytes of APS patients with thrombosis but decreased as compared with monocytes from APS without thrombosis [32]. This may suggest that the downregulation and upregulation of RHOA may be directly associated to the thrombotic state. Significant downregulation of RHOA in our study may indicate the association of this marker to mechanisms of APS other than thrombosis. The other 3 genes have never been reported in any APS study. However, the role of KIT was stressed in the development of placenta tissue in-utero. mRNA expression of KIT detected at the stages of preimplantation and placenta development is suggestive of its role in promoting proliferation and differentiation of the placenta cells [33]. Suppression of KIT in our findings may indicate placental insufficiency which could be one of the pathogenic mechanisms of APS.

From the results of RET and DUSP10 qPCR quantification, the expressions were 3 times higher in APS compared to normal sample. DUSP10 functions by inactivating their target kinases through dephosphorylating both phosphoserine/threonine and phosphotyrosine residue. DUSP10 negatively regulate members of MAP kinase superfamily which are associated with cellular proliferation and differentiation. The upregulated DUSP10 might have an association to downregulated level of KIT as DUSP10 functions to reduce activated tyrosine kinases which further results in desensitization of the kinase receptor. RET is a member of the cadherin superfamily, which is encoded for receptor tyrosine kinases, a cell-surface molecule that transduces signal for cell growth and differentiation [34]. RET mutation was previously reported to increase angiogenesis in thyroid carcinoma [35, 36]. APS placenta was reported to have poor blood circulation due to infarction [37]. In response to infarction, APS placenta might have increased vascular formation as a compensation mechanism. Our results on RET expression may reflect the hypothesis where RET may act as an indicator of increased angiogenesis. A study by Caroll et al found that the treatment of aPLs on trophoblast cells altered the trophoblasts’ angiogenic factors, where the anti-β2GPI-exposed trophoblast produced higher VEGF and PIGF [28].

miR-525-5p was validated by qPCR and showed significant downregulation in APS sample with large fold changes (Table 17). miR-525-5p was predicted to target CD19 and MAPKAPK2. However, only MAPKAPK2 expression was upregulated and in correlation to the reduced expressions of miR-525-5p but the data was not significant. CD19 may not be the true target of miR-525-5p or it is also a target of other upregulated miRNAs, as one gene can be controlled by more than one miRNA and one miRNA can regulate multiple genes. Hromadnikova et al reported that the downregulation of some C19MC miRNAs including miR-525-5p was found to be a common phenomenon shared between placenta tissue from gestational hypertension, preeclampsia and foetal growth restriction. Further analysis on predicted miR-525-5p’s target genes using miRWalk database revealed other genes than CD19 and MAPKAPK2 [38].

By using pathway analysis, BLK was identified as the downregulated miRNA-targeted gene (Fig 5). BLK is known to be involved in B-cell activation. BLK is targeted by other miRNAs which were not validated by qPCR in this study, miR-564 and miRNA-4497. Therefore, we could not make a correlation between miRNAs related to BLK and their regulations on BLK in Obstetric APS. BLK gene encodes for a tyrosine kinase. This tyrosine kinase is involved in the regulation of B-cell activation and may influence the proliferation and differentiation of B-cells [39]. Studies showed that BLK expression is associated with primary APS [40] and SLE [41, 42]. Presence of SLE in antiphospholipid syndrome is categorised as secondary APS. In addition, association of BLK with other rheumatic diseases such as rheumatoid arthritis (RA) and multiple sclerosis [41] has also been reported. This suggests that BLK could be the vulnerable gene in Obstetric APS. Hence, regulation of BLK by miRNAs should be further investigated.

Another downregulated miRNA-targeted gene, MYH15, has never been reported to be involved in antiphospholipid syndrome or other rheumatic diseases. Recent study has suggested the role of MYH15 in Amyotrophic lateral sclerosis (ALS), a fatal neurological disorder characterized by progressive muscular atrophy and respiratory failure [43]. The reduced expression of MYH15 in APS sample by qPCR was also not in concordant to its downregulated miRNAs. However, MYH15 is involved in inflammatory signalling cascade and further investigation could clarify its regulation in APS.

This study therefore indicates the useful strategy of miRNA to narrow down the cohort of candidate transcriptomes that can be mapped to either biological processes or on the genes from pathways of interest in relation to the cell type or disease under investigation. Most notably, this is the first study to our knowledge, to investigate the profiling of miRNA from patient’s tissue sample in obstetric APS. Our results provide compelling evidence for the involvement of certain genes to explain the molecular mechanism of APS. However, some limitations are worth noting. Although our hypotheses were supported statistically, the various experimental settings, normalization application and platforms that may result in inconsistent outcomes of the differential expression of miRNA in the placenta tissues required strict monitoring. However, we have kept these variables to a minimum by adhering to protocols, optimizing each step before actual test samples are run and running duplicates for each test run. The inclusion criteria could have also been made to be more homogeneous in terms of clinical criteria or gestational week among pregnant women of APS. Future work should therefore include follow-up work designed to evaluate these preliminary finding at a larger scale. Future multicentre studies should be carried out to obtain larger sample size which best reflect APS in obstetrics. Further work should also include the characterization of significantly upregulated and downregulated genes identified in the current work at the protein level using RNA interference strategies.

## 5.0 Conclusion

We have identified the genetic expression of VE-cadherin, RHOA, RET, KIT and DUSP10 in VEGFA-VEGFR signalling pathway in placenta tissue of obstetric APS patients as compared to normal placenta. This finding suggests the inducing effect of antiphospholipid antibodies on miRNAs which control the regulation of genes in vascular-associated pathways. The data also demonstrated that not only miRNAs expressions were induced but some miRNAs were supressed. Inhibition of certain miRNAs may cause an overexpression of its targeted genes. It would be interesting to further characterise other miRNAs and its targeted genes to illustrate miRNA-associated mechanism of APS. miR-525-5p for example was greatly downregulated in APS thus, its associated genes would require further investigation by functional analysis. The relationship between miRNAs and its new target genes in placenta tissue can also be established as a novel database on miRNA as the targeted genes are mainly based on cancer studies.

## Acknowledgment

This study has been made possible by funding from the Ministry of Higher Education Malaysia, under the Exploratory Research Grant Scheme (ERGS) ERGS/1/2013/SKK06/USIM/02/2 and registered and approved by the Medical Research Ethical Committee of Malaysia, NMRR-14-1234-22006 and was conducted in compliance with the study protocol and Clinical Research Centre (CRC)’s standard operating procedures.

## Reference

1. Wilson WA, Gharavi AE, Koike T, Lockshin MD, Branch DW, Piette J-C, et al. International consensus statement on preliminary classification criteria for definite antiphospholipid syndrome: Report of an International workshop. 1999;42(7):1309–11.

2. Alijotas-Reig J, Ferrer-Oliveras R. The European Registry on Obstetric Antiphospholipid Syndrome (EUROAPS): A preliminary first year report. Lupus. 2012;21(7):766–8.

3. Ruiz-Irastorza G, Crowther M, Branch W, Khamashta MA. Antiphospholipid syndrome. The Lancet [Internet]. 2010 2010/10/30/; 376(9751):[1498–509 pp.]. Available from: http://www.sciencedirect.com/science/article/pii/S014067361060709X.

4. Out HJ, Kooijman CD, Bruinse HW, Derksen RH. Histopathological findings in placentae from patients with intra-uterine fetal death and anti-phospholipid antibodies. European Journal of Obstetrics & Gynecology and Reproductive Biology [Internet]. 1991 14/06/2019; 41(3):[179–86 pp.].

5. Choi S-Y, Yun J, Lee O-J, Han H-S, Yeo M-K, Lee M-A, et al. MicroRNA expression profiles in placenta with severe preeclampsia using a PNA-based microarray. Placenta [Internet]. 2013; 34(9):[799–804 pp.].

6. Sioud M, Cekaite L. Profiling of miRNA Expression and Prediction of Target Genes. In: Sioud M, editor. RNA Therapeutics: Function, Design, and Delivery. Totowa, NJ: Humana Press; 2010. p. 255–69.

7. Prieto DMM, Markert UR. MicroRNAs in pregnancy. Journal of Reproductive Immunology. 2011;88(2):106–11.

8. Bidarimath M, Khalaj K, Wessels JM, Tayade C. MicroRNAs, immune cells and pregnancy. Cell Mol Immunol. 2014;11(6):538–47.

9. Zhu Y, Lu H, Huo Z, Ma Z, Dang J, Dang W, et al. MicroRNA-16 inhibits feto-maternal angiogenesis and causes recurrent spontaneous abortion by targeting vascular endothelial growth factor. Scientific Reports [Internet]. 2016 14/06/2019; 6:[35536 p.]. Available from: https://doi.org/10.1038/srep35536.

10. Jackson A, Linsley PS. The therapeutic potential of microRNA modulation. Discovery medicine. 2010;9(47):311–8.

11. Mi H, Thomas P. PANTHER pathway: an ontology-based pathway database coupled with data analysis tools. Methods in molecular biology (Clifton, NJ). 2009;563:123–40.

12. Fabregat A, Jupe S, Matthews L, Sidiropoulos K, Gillespie M, Garapati P, et al. The Reactome Pathway Knowledgebase. Nucleic Acids Res. 2018;46(D1):D649–D55.

13. Tüfekci KU, Öner MG, Meuwissen RLJ, Genç Ş. The Role of MicroRNAs in Human Diseases. In: Yousef M, Allmer J, editors. miRNomics: MicroRNA Biology and Computational Analysis. Totowa, NJ: Humana Press; 2014. p. 33–50.

14. Teruel R, PÉRez-SÁNchez C, Corral J, Herranz MT, PÉRez-Andreu V, Saiz E, et al. Identification of miRNAs as potential modulators of tissue factor expression in patients with systemic lupus erythematosus and antiphospholipid syndrome. Journal of Thrombosis and Haemostasis. 2011;9(10):1985–92.

15. Deng Y, Zhao J, Sakurai D, Kaufman KM, Edberg JC, Kimberly RP, et al. MicroRNA-3148 Modulates Allelic Expression of Toll-Like Receptor 7 Variant Associated with Systemic Lupus Erythematosus. PLOS Genetics [Internet]. 2013; 9(2):[e1003336 p.]. Available from: https://doi.org/10.1371/journal.pgen.1003336.

16. Shi C, Xu X. MiR-670-5p induces cell proliferation in hepatocellular carcinoma by targeting PROX1. Biomedicine & Pharmacotherapy [Internet]. 2016; 77:[20–6 pp.].

17. Tammela T, Alitalo K. Lymphangiogenesis: molecular mechanisms and future promise. Cell [Internet]. 2010; 140(4):[460–76 pp.].

18. Wang Y. Vascular biology of the placenta: Morgan & Claypool Life Sciences; 2010. 1–98 p.

19. Krissansen GW, Yang Y, McQueen FM, Leung E, Peek D, Chan YC, et al. Overexpression of miR-595 and miR-1246 in the sera of patients with active forms of inflammatory bowel disease. Inflammatory bowel diseases [Internet]. 2015; 21(3):[520–30 pp.].

20. Madhavan B, Yue S, Galli U, Rana S, Gross W, Müller M, et al. Combined evaluation of a panel of protein and miRNA serum-exosome biomarkers for pancreatic cancer diagnosis increases sensitivity and specificity. International journal of cancer [Internet]. 2015; 136(11):[2616–27 pp.].

21. Choi SY, Yun J, Lee OJ, Han HS, Yeo MK, Lee MA, et al. MicroRNA expression profiles in placenta with severe preeclampsia using a PNA-based microarray. Placenta. 2013;34(9):799–804.

22. Enquobahrie DA, Abetew DF, Sorensen TK, Willoughby D, Chidambaram K, Williams MA. Placental microRNA expression in pregnancies complicated by preeclampsia. American Journal of Obstetrics & Gynecology. 2011;204(2):178.e12–.e21.

23. Ventura W, Koide K, Hori K, Yotsumoto J, Sekizawa A, Saito H, et al. Placental expression of microRNA-17 and −19b is down-regulated in early pregnancy loss. European Journal of Obstetrics & Gynecology and Reproductive Biology. 2013;169(1):28–32.

24. Johnson KE, Wilgus TA. Vascular Endothelial Growth Factor and Angiogenesis in the Regulation of Cutaneous Wound Repair. Adv Wound Care (New Rochelle). 2014;3(10):647–61.

25. Di Simone N, Di Nicuolo F, D’Ippolito S, Castellani R, Tersigni C, Caruso A, et al. Antiphospholipid antibodies affect human endometrial angiogenesis. Biol Reprod. 2010;83(2):212–9.

26. Mulla MJ, Myrtolli K, Brosens JJ, Chamley LW, Kwak-Kim JY, Paidas MJ, et al. Antiphospholipid antibodies limit trophoblast migration by reducing IL-6 production and STAT3 activity. American journal of reproductive immunology (New York, NY: 1989). 2010;63(5):339–48.

27. Ichikawa G, Yamamoto T, Chishima F, Nakamura A, Kuno S, Murase T, et al. Effects of anti-beta2-glycoprotein I antibody on PlGF, VEGF and sVEGFR1 production from cultured choriocarcinoma cell line. The journal of obstetrics and gynaecology research. 2011;37(8):1076–83.

28. Carroll TY, Mulla MJ, Han CS, Brosens JJ, Chamley LW, Giles I, et al. Modulation of trophoblast angiogenic factor secretion by antiphospholipid antibodies is not reversed by heparin. American journal of reproductive immunology (New York, NY: 1989). 2011;66(4):286–96.

29. Di Simone N, Castellani R, Caliandro D, Caruso A. Antiphospholid antibodies regulate the expression of trophoblast cell adhesion molecules. Fertility and Sterility. 2002;77(4):805–11.

30. Damsky CH, Librach C, Lim KH, Fitzgerald ML, McMaster MT, Janatpour M, et al. Integrin switching regulates normal trophoblast invasion. Development (Cambridge, England). 1994;120(12):3657–66.

31. Rolfe BE, Worth NF, World CJ, Campbell JH, Campbell GR. Rho and vascular disease. Atherosclerosis. 2005; 183(1):1–16.

32. Lopez-Pedrera C, Cuadrado MJ, Herandez V, Buendia P, Aguirre MA, Barbarroja N, et al. Proteomic analysis in monocytes of antiphospholipid syndrome patients: deregulation of proteins related to the development of thrombosis. Arthritis and rheumatism. 2008;58(9):2835–44.

33. Horie K, Fujita J, Takakura K, Kanzaki H, Kaneko Y, Iwai M, et al. Expression of C-Kit Protein during Placental Development1. Biology of Reproduction. 1992;47(4):614–20.

34. de Groot JWB, Links TP, Plukker JTM, Lips CJM, Hofstra RMW. RET as a Diagnostic and Therapeutic Target in Sporadic and Hereditary Endocrine Tumors. Endocrine Reviews. 2006;27(5):535–60.

35. Dunna NR, Kandula V, Girdhar A, Pudutha A, Hussain T, Bandaru S, et al. High Affinity Pharmacological Profiling of Dual Inhibitors Targeting RET and VEGFR2 in Inhibition of Kinase and Angiogeneis Events in Medullary Thyroid Carcinoma. Asian Pacific journal of cancer prevention: APJCP. 2015;16(16):7089–95.

36. Verrienti A, Tallini G, Colato C, Boichard A, Checquolo S, Pecce V, et al. RET mutation and increased angiogenesis in medullary thyroid carcinomas. Endocrine-related cancer. 2016;23(8):665–76.

37. Van Horn JT, Craven C, Ward K, Branch DW, Silver RM. Histologic features of placentas and abortion specimens from women with antiphospholipid and antiphospholipid-like syndromes. Placenta. 2004;25(7):642–8.

38. Hromadnikova I, Kotlabova K, Ondrackova M, Pirkova P, Kestlerova A, Novotna V, et al. Expression profile of C19MC microRNAs in placental tissue in pregnancy-related complications. DNA Cell Biol. 2015;34(6):437–57.

39. Zhang H, Wang L, Huang Y, Zhuang C, Zhao G, Liu R, et al. Influence of BLK polymorphisms on the risk of rheumatoid arthritis. Molecular Biology Reports. 2012;39(11):9965–70.

40. Yin H, Borghi MO, Delgado-Vega AM, Tincani A, Meroni PL, Alarcon-Riquelme ME. Association of STAT4 and BLK, but not BANK1 or IRF5, with primary antiphospholipid syndrome. Arthritis and rheumatism. 2009;60(8):2468–71.

41. Tsuchiya N, Ito I, Kawasaki A. Association of IRF5, STAT4 and BLK with systemic lupus erythematosus and other rheumatic diseases. Nihon Rinsho Men’eki Gakkai kaishi = Japanese journal of clinical immunology. 2010;33(2):57–65.

42. Castillejo-López C, Delgado-Vega AM, Wojcik J, Kozyrev SV, Thavathiru E, Wu Y-Y, et al. Genetic and Physical Interaction of the B-Cell SLE-Associated Genes BANK1 and BLK. Annals of the Rheumatic Diseases. 2012;71(1):136–42.

43. Kim H, Lim J, Bao H, Jiao B, Canon SM, Epstein MP, et al. Rare variants in MYH15 modify amyotrophic lateral sclerosis risk. Human Molecular Genetics. 2019.

